# Quantifying mechanical loading and elastic strain energy of the human Achilles tendon during walking and running

**DOI:** 10.1101/2020.09.05.284182

**Authors:** Mohamadreza Kharazi, Sebastian Bohm, Christos Theodorakis, Falk Mersmann, Adamantios Arampatzis

## Abstract

The purpose of the current study was to assess Achilles tendon (AT) mechanical loading and strain energy during locomotion using a new in vivo approach for measuring AT length that considers the AT curve-path shape. Eleven participants walked at 1.4 m/s and ran at 2.5 m/s and 3.5 m/s on a treadmill. AT length, defined as the distance between its origin at the gastrocnemius medialis myotendinous junction (MTJ) and the calcaneal insertion, was determined experimentally by integrating kinematics and ultrasound analysis. Small foil markers were placed on the skin covering the AT path from the origin to the insertion, and the MTJ, tracked using ultrasonography, was projected to the reconstructed skin to account for their misalignment. Skin-to-bone displacements were assessed during a passive rotation (5 °/s) of the ankle joint and considered in the calculation of AT length. Force and strain energy of the AT during locomotion were calculated by fitting a quadratic function to the experimentally measured tendon force-length curve obtained from maximum voluntary isometric contractions. Maximum AT strain and force were affected by speed (p<0.05, ranging from 4.0 to 4.9% strain and 1.989 to 2.556 kN), yet insufficient in magnitude to be considered an effective stimulus for tendon adaptation. Further, we found a recoil of elastic strain energy at the beginning of the stance phase of running (70-77 ms after touch down) between 1.7 ±0.6 and 1.9 ±1.1 J, which might be functionally relevant for running efficiency.

**Summary statement:** A new accurate in vivo approach to assess Achilles tendon strain, force and strain energy during locomotion.

## Introduction

Tendons cannot generate force actively, yet their elastic behavior upon loading influences the function of the muscle-tendon unit during locomotion. The time course of the human Achilles tendon (AT) length during functional tasks as walking, running, jumping, or cycling is important to understand the interaction between muscle and tendon (Lichtwark and Wilson, 2005, Dick and Wakeling, 2017, Kümmel et al., 2018), to assess tendon loading (Dick et al., 2016) and to examine tendon elastic strain energy (Ishikawa et al., 2005, Lai et al., 2014, Monte et al., 2020). The reported maximum strain values of the AT during running are between 4.6% and 9.0% (Lichtwark et al., 2007, Lai et al., 2015, Lai et al., 2018) and between 4.0 to 4.3% during walking (Lichtwark et al., 2007, Ishikawa et al., 2005). Corresponding AT force values ranged from 3.06 to 4.64 kN during running (Almonroeder et al., 2013, Werkhausen et al., 2019) and about 2.63 kN during walking (Giddings et al., 2000). These results suggest substantial mechanical loading of the AT during human locomotion. The AT can adapt to external mechanical loading by increasing its stiffness, elastic modulus and size (Bohm et al., 2015, Wiesinger et al., 2016). Loading-induced alteration of tendon properties is a biological mechanism to maintain the functional integrity of the muscle-tendon unit and to keep tendon mechanical loading in a physiological range during functional tasks (Wang et al., 2013, Arnoczky et al., 2002). Repetitive loading of the AT with a strain magnitude between 4.5 and 6.5% has been evidenced as an effective mechanical stimulus for the improvement of AT mechanical properties (Arampatzis et al., 2007a, Arampatzis et al., 2010, Bohm et al., 2014). This tendon strain range is commonly reached at about 90% of a voluntary maximum isometric contraction of the adjacent muscle, which results in AT forces between 2.34 and 3.69 kN (Arampatzis et al., 2007; 2010; Bohm et al., 2014).

Although the reports of AT strain and force during running indicate sufficient AT loading for the initiation of adaptive alterations in tendon properties (Lai et al., 2015, Lai et al., 2018, Lichtwark et al., 2007, Almonroeder et al., 2013) most studies, which compared the AT mechanical properties between runners and untrained controls were not able to detect any differences between the two groups (Karamanidis and Arampatzis, 2005, Arampatzis et al., 2007b, Kubo et al., 2010, Wiesinger et al., 2016). Furthermore, in the longitudinal study of Hansen et al. (2003), no significant changes in the mechanical properties of AT were observed after nine months (78 sessions) of running training (Hansen et al., 2003). This discrepancy (estimates of loading vs. lack of adaptive response) might be the result of the methodological approaches used for the assessment of AT strain during locomotion in vivo. The in vivo approaches used for assessing AT length during functional tasks either calculated AT length using a simple planimetric model (Lichtwark et al., 2007, Lai et al., 2015) or did not consider the concave curvature of the AT in their measurements (Lichtwark and Wilson, 2005, Lichtwark and Wilson, 2006, Dick et al., 2016). There is evidence that considering the AT as a straight line between calcaneus (insertion) and gastrocnemius medialis (GM) myotendinous junction (MTJ, origin) results in an overestimation of the AT length and substantial errors (up to 78%) of the AT length changes (Fukutani et al., 2014).

The instantaneous curved length of the AT can be obtained by using a line of reflective markers attached to the skin from the tuber calcanei to the MTJ of the GM (De Monte et al., 2006, Arampatzis et al., 2008). Although this method has been validated by comparing the outcomes with accurate measurements of AT length from magnet resonance imaging (Fukutani, 2014), it has not been applied for the assessment of AT length during locomotion yet. However, during dynamic functional tasks, two more issues may introduce errors in such AT length measurements. First, the original position of the GM MTJ is not aligned with the reflective markers attached on the skin surface and, therefore, for an accurate AT length measurement, the identified position of the MTJ should be projected to the skin (Fig. 1A). Second, potential displacement of the skin to the bone underneath the calcaneus marker that defines the AT insertion can also introduce artifacts in the AT length measurement (Lichtwark and Wilson, 2005).

**Figure 1.**
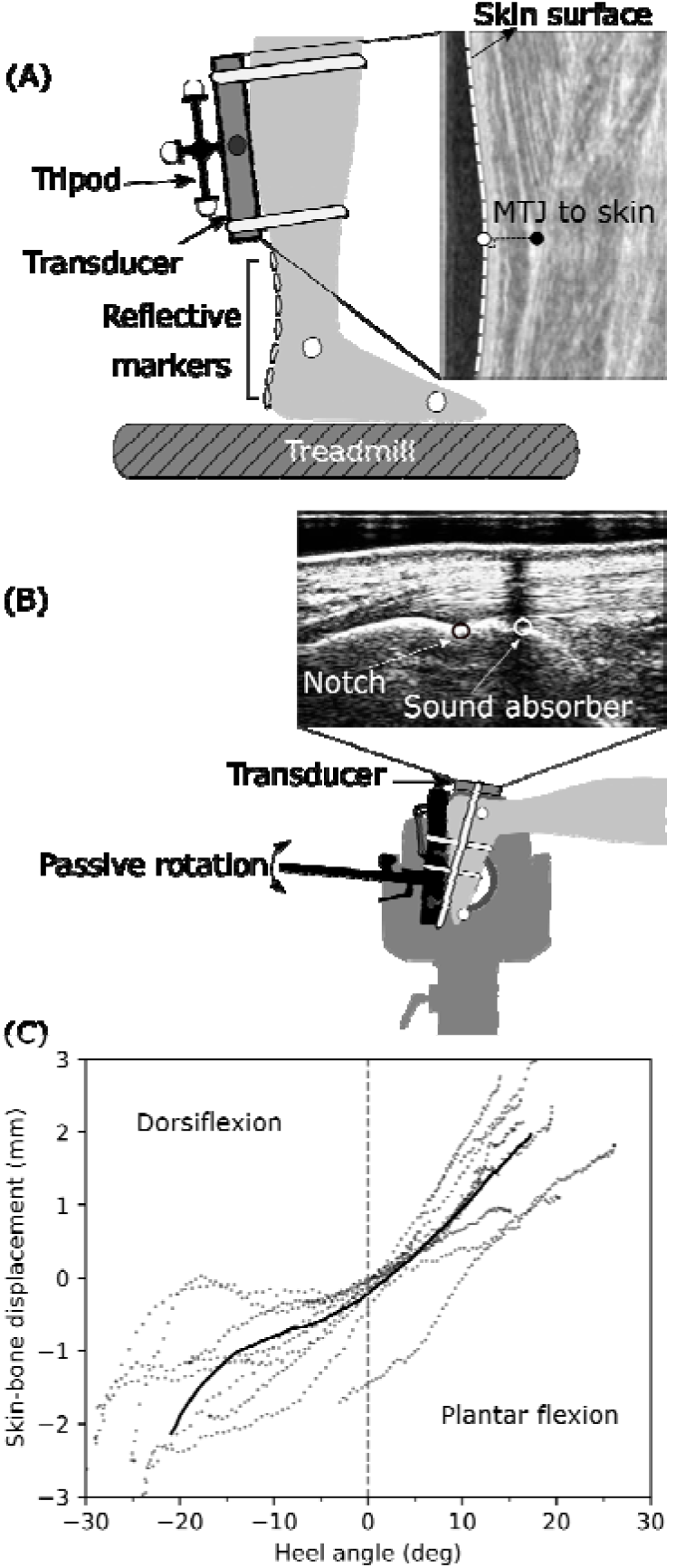
(A) Experimental setup for the determination of Achilles tendon (AT) length during gait. A) Reflective foil markers on the skin were used to reconstruct the curve-path shape of the AT. The position of the gastrocnemius medialis myotendinous junction was projected to the skin surface, and the coordinates of the ultrasound images were transformed to the global coordinate system using a marker tripod attached to the ultrasound probe. (B) A three-centimeter ultrasound probe was placed on top of the calcaneus bone and a sound absorptive marker in-between to measure the movement of the calcaneus bone (notch) with regard to the skin (absorber marker) as a function of the heel to shank angle. (C) Average (solid line) and individual (filled dots) data of skin-to-bone displacements vs. heel angle. Positive heel angles represent plantar flexion, and negative ones are dorsiflexion.

The elastic strain energy recoil of the AT during the propulsion phase of walking and running is a well-known mechanism within the muscle-tendon unit, which increases the efficiency of muscle power output (Lai et al., 2014, Ishikawa et al., 2005, Monte et al., 2020). The contribution of the elastic strain energy recoil to the positive work generated by the muscle-tendon unit is greater compared to the work produced by the muscle fascicles alone (Lai et al., 2014, Ishikawa et al., 2005, Monte et al., 2020). However, there are also indications of AT elastic strain energy recoil during the early stance phase of running (Komi, 1990), which is not well understood until now. Komi (1990), using an implanted transducer around the AT, measured in vivo AT forces during running and found an apparent decrease of AT force after heel contact, particularly in rearfoot running, indicating an energy recoil of the AT directly after touchdown. Considering that in this phase, the fascicles of the triceps surae muscles actively shorten (Lichtwark et al., 2007, Lai et al., 2018, Bohm et al., 2019), it can be argued that this elastic strain energy recoil from the AT is not absorbed by the contractile elements and, thus, might be an additional important source of energy for human running.

The aim of the current study was to assess the mechanical loading and strain energy of the AT during walking and running. For this purpose, we measured the AT length during walking and running using a new in vivo approach, which considers the tendon curve-path shape using skin markers, taking into account the projection of the MTJ to the skin surface as well as potential movements of the skin relative to the calcaneus bone. Tendon force and strain energy were then calculated based on an experimentally determined tendon force-elongation relationship. We hypothesized a relevant contribution of MTJ projection to the skin as well as skin-bone displacement on the AT length. Further, we expected lower levels of tendon strain during locomotion, as previously reported, insufficient in magnitude to serve as a stimulus for the adaptation of AT mechanical properties. Finally, we hypothesized a functional relevant recoil of tendon elastic strain energy to the body at the beginning of the stance phase during running.

## Methods

### Experimental design

Eleven adults (one female) with an average height of 177 ± 6 cm, the body mass of 74 ± 9 kg and the age of 29 ± 3 years participated in this study. All participants gave written informed consent to the experimental procedure, which was approved by the ethics committee of the Humboldt-Universität zu Berlin (HU-KSBF-EK_2018_0005) and followed the standards of the Declaration of Helsinki. The participants were in full health and none of them reported neuromuscular or skeletal impairments in the past year. After a familiarization phase, the participants walked (1.4 m/s) and ran (2.5 m/s and 3.5 m/s) on a treadmill (Daum electronic, ergo_run premium8, Fürth, Germany) whereby AT length was determined experimentally by integrating kinematics and ultrasound analysis. AT length was defined as the distance between the origin (i.e., the most distal junction) of the GM and the insertion on the tuber calcanei.

During gaits, marker-based motion capture was used to track the position of the insertion point, represented by a marker that was placed over the tuber calcanei as well as joint kinematics (Fig. 1A). The potential skin-to-bone displacement was considered while the foot was passively rotated (5°/s) throughout the full range of motion from plantar flexion to dorsiflexion by means of ultrasonography (Fig. 1B). Ultrasound was used for the detection of the MTJ, and the position of the transducer was tracked by the motion capture system using a mounted marker tripod (Fig. 1A). To assess the curved path of the AT, small foil markers were placed on the skin covering the AT path on the line from the defined origin to the insertion (Fig. 1A). The MTJ position was then projected to the skin (Fig. 1A) and mapped to the global coordinate system to be able to assess AT length during gait (i.e., from projected MTJ over the curved foil marker path to insertion). Force and strain energy of AT during locomotion were assessed by fitting a quadratic function to the experimentally measured force-length curve of the AT based on maximum voluntary isometric contractions (MVC) for each participant. The coefficients of the quadratic function were applied to the AT elongation values obtained during locomotion. The strain energy of the AT tendon during walking and running was calculated by integrating tendon force over tendon elongation.

### Kinematics and gait cycle determination

Kinematic data of the right leg were assessed employing six reflective markers (14 mm in diameter) that were placed on the tip of the toe, medial and lateral epicondyle, on a line from greater-trochanter to lateral epicondyle as well as medial and lateral malleolus. 3D trajectories of all markers were captured in real-time with 14 Vicon (Version 1.7.1, Vicon Motion Systems, Oxford, UK) cameras (4x MX T20, 2x MX-T20-S, 6x MX F20, 2x MX F40, 250 Hz). A fourth-order low pass and zero-phase shift Butterworth filter with a cut off frequency of 7 Hz was applied to the raw marker trajectories. A one-minute warm-up and familiarization phase with the treadmill in each of the three speeds was considered prior to the captured trials, which included at least twelve stride cycles for each speed. The touchdown of the foot during walking was determined as the instant minimal vertical position of the heel marker (Fellin et al., 2010) and during running as the first peak of the knee joint angle (i.e., extension) (Dingwell et al., 2001). The take-off was defined as the reversal of the anterior-posterior velocity of the toe marker during walking and as the second peak of the knee joint angle during running (Alvim et al., 2015).

### Measurement of the AT length during gait

The point of insertion of the AT, defined at the notch of the tuber calcanei of the calcaneus bone (Kongsgaard et al., 2011), was detected in sagittal plane ultrasound scans. The origin of the AT was determined as the most distal position of the GM MTJ. A T-shaped 60 mm ultrasound transducer (Aloka UST-5713T, Hitachi Prosound, alpha 7, Japan) operating at 146 Hz was fixed over the MTJ of the AT with a customized flexible plastic cast. A gel pad was used to account for surface unevenness. The ultrasound device was time-synchronized with the motion capture system using a manual analog trigger signal. The curved path of the AT was elaborated by placing reflective foil markers on the skin that directly covers the AT (De Monte et al., 2006, Arampatzis et al., 2008). Depending on the position of the MTJ on the shank length, a varying number (i.e., from 5 to 9) of reflective plane foil markers with 5 millimeters in diameter and 20 millimeters interval gap were placed on the path of AT from the defined insertion point to the last possible position below the cast. Note that the ultrasound probe was then oriented in extension to this line. The length of the curved path was calculated as the sum of the vectors, which was defined by the position of two consecutive foil markers.

An image-based tracking algorithm was developed to determine the position of the MTJ from the ultrasound videos. The procedure included a multi-updating template-matching technique with 33 manually defined templates. These templates are rectangular windows covering the area around the MTJ and were defined in the first step cycle, distributed equally in time throughout the stance and swing phase. The created templates then served to detect the MTJ during the subsequent steps (10 step cycles of the right leg on average). More specifically, the ultrasound images were first cropped to a region of interest and were convolved with a gaussian (3*3 kernel size) filter to reduce speckles. By sweeping the each defined template on the cropped image, the maximum value of normalized 2D cross-correlation was detected as the position of MTJ. Each template auto-updates itself under specific criteria until the next manually defined template occurs. The criteria of the auto-updating template were defined as Eqn 1 and Eqn 2:

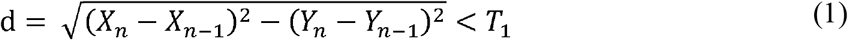

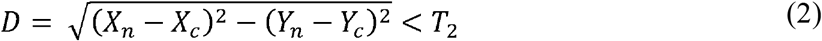

Where ***X_n_*** and ***Y_n_*** are the coordinates of the best-matched position (MTJ coordination) in the current frame, ***X_n−1_*** and ***Y_n−1_*** are the best-matched position in the previous frame, ***X_c_*** and ***Y_c_*** are the coordination of best-matched position in the last accepted frame (i.e., the last frame that template was auto-updated), ***T*_1_** and ***T*_2_** are the thresholds, which were defined in pixel (in our case was 5 and 10 pixels respectively), ***d*** is the Euclidean distance between the best-matched position of the current frame and the previous frame, and **D** is the Euclidean distance between the current frame and the last accepted frame. If ***d*** and ***D*** were below the thresholds, then the updated template was accepted and used for the next frame. If the criteria were not fulfilled, the last accepted template was used for the next frame. Visual inspection of all frames was conducted afterward to verify the automatic tracking results, and corrections were made if necessary. When an inappropriately tracked MTJ position was deleted, the gap was filled by linear interpolation. In case the interpolation was inacceptable, the position of MTJ was defined manually.

A custom 3D-printed marker tripod was fixed to the ultrasound transducer. The calibration of the US image with respect to the tripod was done by digitizing the four corners of the transducer’s protective front layer, and a coordinate system was defined (P) on the center-left side of the transducer. The gap between the left edge side of the protective front layer to the first piezoelectric sensor was determined by subtracting the width of the plate from the width of the ultrasound image divided by two. This gap size was then verified with an x-ray image of the transducer. The origin of the coordinate system (t) on the transducer (defined by a mounted tripod) was then adjusted accordingly. Another coordinate system (2D) was determined with the origin located at the first pixel of the ultrasound image (U). To map the MTJ position from the ultrasound image to the global coordinate system, a global transformation matrix 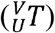 was calculated by multiplication of the three transformation matrixes (Eqn 3) (Craig, 2005, Rasske and Franz, 2018).

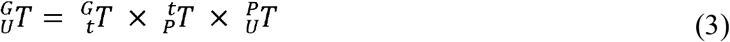

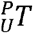 is the transformation matrix to transform the coordinate systems from the ultrasound image [U] to front face of the transducer [P], 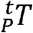 is the transformation matrix for the transformation of the transducer to tripod [t] and 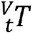 transfer the tripod coordination to the global coordination system [G] (i.e., defined by the motion capture system). In this way, the detected MTJ position from the ultrasound images could be projected to the global coordination system (Eqn 4):

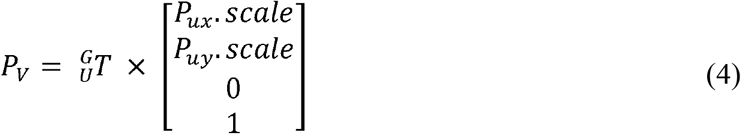

where *P_v_* is the transformed MTJ position to the global coordinate system, *P_ux_* and *P_uy_* are the detected MTJ position in the vertical and horizontal direction, respectively, and *scale* is the pixel to millimeter scale-factor for the ultrasound image. For the projection of the MTJ to the skin surface, a double threshold-to-intensity gradient of the image (Canny, 1987) was applied at a selected range of interest of the image. Then a spline curve was fitted to the detected edges of the skin surface, and the position of the MTJ was transferred to the fitted curve via the shortest distance in each frame.

The kinematic and ultrasound recordings obtained to account for potential displacements of the reflective calcaneus marker above the defined insertion point (notch at tuber calcanei) were captured separately due to spatial constraints around the ankle. In the first step, the participants lay prone on a dynamometer bench (Biodex Medical, Syst. 3, Inc., Shirley, NY) while their foot was fixed to the dynamometer’s footplate. Three trials were recorded while the foot was rotated passively by the dynamometer (5°/s) from 30° plantar flexion to the individual maximum dorsiflexion angle (Fig. 1B). A three-centimeter ultrasound transducer (My Lab60, Esaote, Genova, Italy, 37 Hz) was attached above the calcaneus bone over the insertion of the AT in a sagittal plane. A sound-absorptive skin-adhesive marker was then placed about 5 millimeters above the identified insertion point. In the ultrasound images, the skin position was represented by the registered shadow line of the sound absorptive marker and the corresponding position of the bone by the notch as a fixed bony landmark. The position of the skin with respect to the bone was tracked throughout the full range of motion from plantar flexion to dorsiflexion. In the second step, the ultrasound probe was removed, and reflective markers were placed precisely on the same positions as during the treadmill session, i.e., epicondyle (lateral and medial), malleoli (lateral and medial), between the first and the second metatarsal and on the defined insertion point (i.e., notch). Then the ankle joint was again passively rotated under the same conditions by the dynamometer. The heel angle (i.e., the angle between the vector between knee joint center and ankle joint center and the vector of heel and ankle joint center) was calculated and matched to the angle given by the dynamometer observed in step one and two. Finally, the differences between skin and bone positions measured in the ultrasound images were converted to millimeters, and a spline curve was fitted to skin-to-bone displacement versus heel angle (Fig. 1C). The individual error model was then used to correct potential skin-to-bone movements as a function of the heel angle during gait. The average maximum displacement of skin relative to bone was 2.3 mm in plantarflexion and 1.9 mm in dorsiflexion during the passive rotation of the ankle.

### Analysis of AT length

To determine the effects of the transverse projection of the MTJ to the skin surface and the displacement of the skin under the calcaneus marker on the AT length, we calculated the (a) AT length: as curve-path under consideration of the skin-to-bone displacement and MTJ-to-skin surface projection, (b) AT length without MTJ projection: curve-path and skin-to-bone displacement, (c) AT length without skin-to-bone displacement: curve-path and projection of MTJ to the skin surface, and (d) AT length without MTJ projection and skin-to-bone displacement: only curve-path. Ten gait cycles were analyzed for each participant and averaged. The contribution of the skin-to-bone displacements, projection of MTJ to the skin surface, and both together were elaborated using the root mean square (RMSE) differences of AT length with and without these considerations through the course of the gait cycle.

### Measurement of muscle electromyographic activity

Surface electromyographic (EMG) data of the tibialis anterior (TA), gastrocnemius medialis (GM) and soleus (Sol) was measured during walking and running using a wireless EMG system (Myon m 320RX, Myon AG, Baar, Switzerland) operating at a sampling frequency of 1000 Hz. The EMG signal was processed using a fourth-order high-pass Butterworth zero-phase filter with a 50 Hz cut-off frequency, a full-wave rectification and then a low-pass zero-phase filter of 20 Hz cut-off frequency was applied. The resultant EMG signal was then normalized to the maximum processed EMG obtained during individual MVCs. The placement of EMG electrodes was based on SENIAM recommendations (Stegeman and Hermens, 2007).

### Assessment of AT strain, force and energy during gait

AT strain during walking and running was calculated by dividing the measured AT length by the AT resting length that was determined during the relaxed state in 20° plantar flexion, where slackness of the AT has been reported previously (De Monte et al., 2006). In order to calculate force and strain energy of the AT during locomotion, a force-elongation relationship of the AT was determined in a separate experiment, combining dynamometry (Biodex-System 3, Biodex Medical System Inc., USA) and ultrasound (My Lab60, Esaote, Genova, Italy, 37 Hz) measurements. Participants performed five isometric plantarflexion ramp MVCs (~5 s gradual increase of force) while their knee was fully extended, and the ankle angle at rest was set to the neutral position (tibia perpendicular to sole). Misalignments of the ankle axis of rotation and dynamometer axis during the MVCs, as well as gravitational and passive moments, were considered through inverse dynamics (Arampatzis et al., 2005). The effect of antagonistic muscle co-activation on the resultant joint moment during the MVCs was taken into account with established means (Mademli et al., 2004). The AT force during contractions was calculated by dividing the ankle joint moment by the lever arm of the AT, which was determined using the tendon-excursion method (An et al., 1984). The changes in the tendon lever arm during the contractions were corrected, as suggested previously (Maganaris et al., 1998). The corresponding elongation of the AT during the five trials for each participant was assessed with a 10 cm linear ultrasound probe fastened over the GM MTJ. The position of the MTJ visualized by ultrasound was tracked in the ultrasonographic images with the semi-automatic tracking algorithm described above. The effects of unavoidable ankle joint rotation during the MVCs that cause displacements of the MTJ on the tendon elongation was corrected by subtracting the MTJ displacement tracked during a passive rotation of the ankle (full range of motion at 5 °/s) (Arampatzis et al., 2008) with respect to the ankle joint angle changes during the MVCs. The tendon force-elongation relationship of the five trials of each participant was averaged to achieve excellent reliability (Schulze et al., 2012). A quadratic function (Eqn 5) was fitted to obtain the individual force-elongation relationship of the AT and then was used to assess AT force during gait:

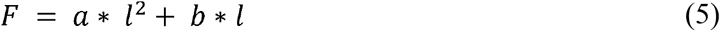

Where **a** and **b** are constants, and ***l*** is the elongation of the AT. The strain energy of the AT during walking and running was calculated by integrating the AT force over the measured AT elongation (we omitted the elongations that were below the resting length) using the Eqn 6.

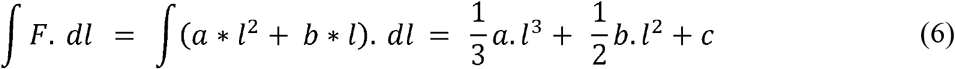

The elastic energy recoil of the AT at the beginning of the stance phase of running was calculated as the difference of the AT strain energy between the touchdown and its first minimum after touchdown.

### Statistics

The validity of the semi-automatic MTJ tracking algorithm was tested by comparison to manual tracking (gold standard). Displacements of the MTJ position in the longitudinal direction were assessed with the parameter variance accounted for (VAF) and the Pearson correlation coefficient in three gait cycles of all gait speeds. To analyze the contribution of each correction method on the AT length, RMSE was calculated. A one-way repeated-measures analysis of variance (ANOVA) was performed to examine the effects of gait speed on AT strain, force, strain energy, EMG activity in GM, Sol and TA and temporal gait parameters such as stance time, swing time and cadence. In case of significant main effects, a pairwise t-test with Benjamini-Hochberg corrected p-values were used for posthoc analysis (adjusted p-values will be reported). The normal distribution of the data was tested with the Shapiro-Wilk test. The level of significance for all statistical tests was set to *α* = 0.05.

## Results

A significant gait speed effect was found in stance time and cadence (*p* < 0.001) but not in swing time (*p* = 0.436, table 1). Comparing semi-automatic versus manual tracking of the MTJ position, the VAF was 93±0.4%, 97±0.1% and 97±0.5%, and the adjusted r^2^ was 0.96±0.02, 0.99±0.008 and 0.98±0.02 (*p* < 0.001) during walking, slow running and fast running, respectively, evidencing high conformity of the developed algorithm with manual tracking (Fig. 2). The AT length over the gait cycle during walking and running is illustrated in Fig. 3. The average RMSE of the skin-to-bone displacement on the AT length was between 0.92 and 1.09 mm, the projection of the MTJ to the skin surface was between 1.9 and 2.1 mm, and both together were between 2.0 and 2.5 mm (table 2 for RMSE).

**Figure 2.**
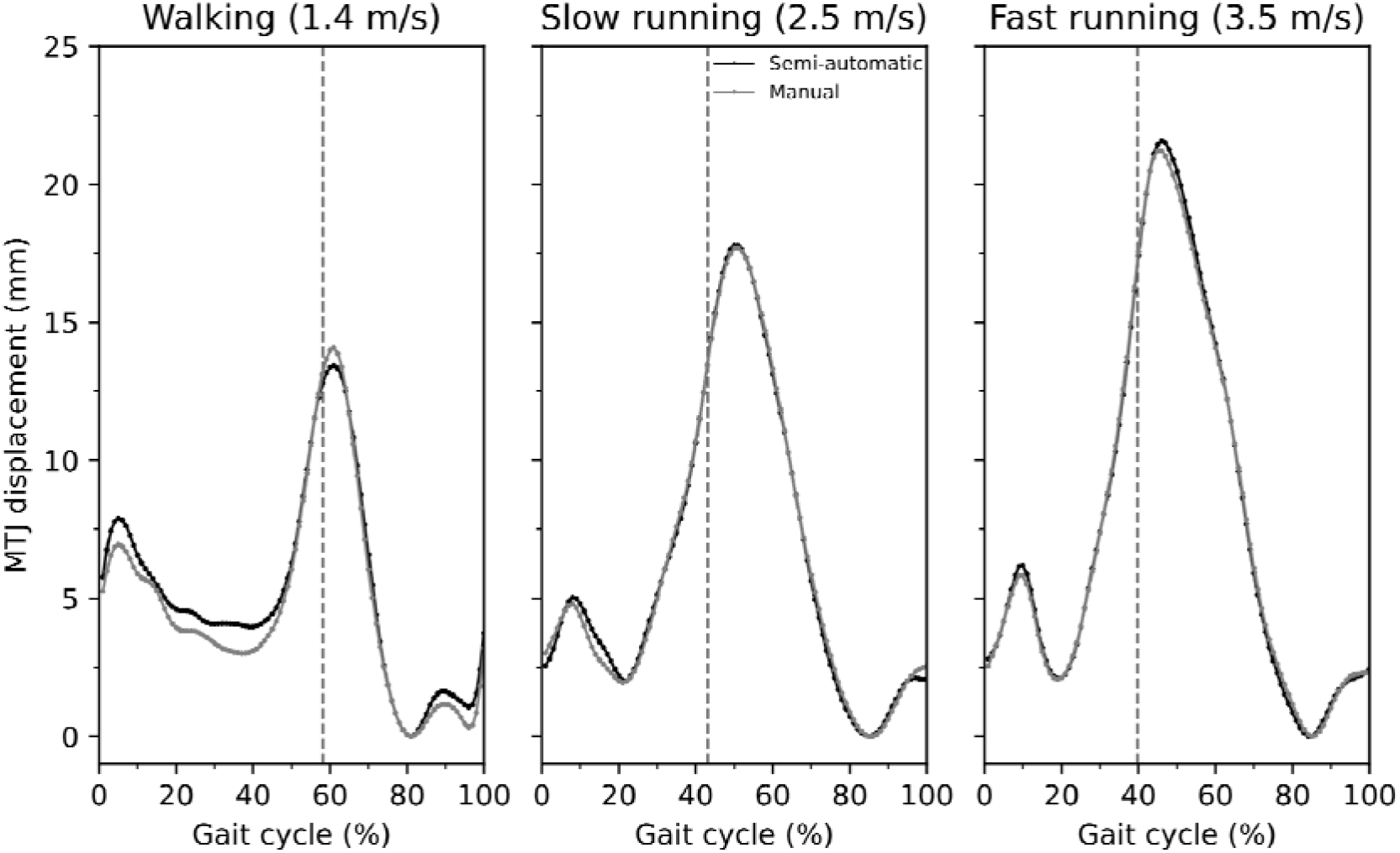
Average values of eleven individuals with ten gait cycle repletion, displacement of gastrocnemius medialis myotendinous junction using the self-developed semi-automatic algorithm in comparison to the manual tracking for walking, slow running, and fast running. The x-axis is normalized to the gait cycle. The gray dashed vertical line separates the contact and swing phase.

**Figure 3.**
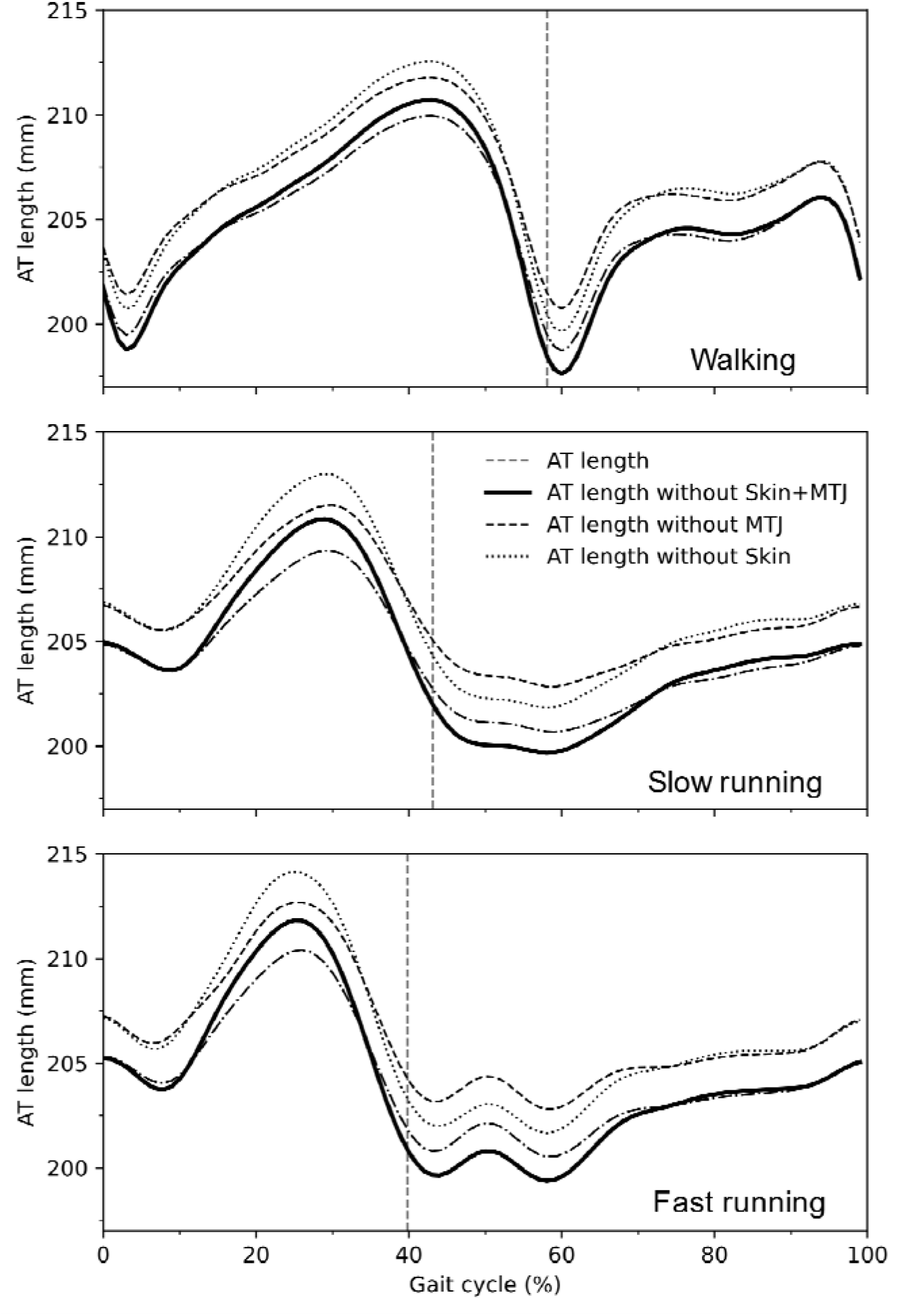
Average values of eleven individuals with ten gait cycle repletion of Achilles tendon (AT) length during walking, slow running (2.5 m/s, middle) and fast running (3.5 m/s, bottom). AT length: length of AT considering curve-path shape, skin-to-bone displacement and projection of myotendinous junction (MTJ) to the skin surface; AT length without skin + MTJ: AT length with only curve-path shape consideration; AT length without MTJ: length of AT considering the curve-path shape and skin-to-bone displacement; AT length without skin: AT length considering curve-path shape and projection of MTJ to the skin surface. The x-axis is normalized to the gait cycle. The gray dashed vertical line separates the contact and swing phase.

**Table 1.** Duration of the stance and swing phases as well as cadence during walking and running (average value ± standard deviation).

|  | Walking (1.4 m/s) | Slow running (2.5 m/s) | Fast running (3.5 m/s) |
| --- | --- | --- | --- |
| Stance (ms) * | 605 ±47 <sup># ^</sup> | 330 ±36 <sup>#</sup> | 279 ±19 |
| Swing (ms) | 437 ±33 | 436 ±45 | 423 ±32 |
| Cadence (gait cycles/s) * | 1.7 ±0.1 <sup># ^</sup> | 2.3 ±0.1 <sup>#</sup> | 2.5 ±0.1 |

\*: Statistically significant gait effect (p<0.05)
<sup>#</sup>: Statistically significant differences (post-hoc analysis) to fast running (p < 0.05)
<sup>^</sup>: Statistically significant differences (post-hoc analysis) to slow running (p < 0.05)

**Table 2.** Contribution of skin-to-bone displacement, projection of MTJ to the skin surface and both two on the AT length as means of root mean square error (average value ± standard deviation).

|  | Walking (1.4 m/s) | Slow running (2.5 m/s) | Fast running (3.5 m/s) |
| --- | --- | --- | --- |
| Skin-Bone (mm) | 0.92 ±0.57 | 1.08 ±0.61 | 1.09 ±0.53 |
| MTJ projection (mm) | 1.90 ±0.90 | 2.00 ±0.80 | 2.10 ±0.80 |
| Skin-Bone +<br>MTJ projection (mm) | 2.00 ±0.80 | 2.20 ±0.80 | 2.50 ±0.80 |

The maximum tendon force and elongation during the isometric MVCs were 5.523±0.552 kN and 14.0±2.5 mm, respectively. The adjusted r^2^ from the force-elongation of the quadratic equation (Eqn 5) was, on average, 0.98 ± 0.01 (*p* < 0.001), and the average values among all participants of the two constants (a and b) were 8.7 and 181 for a and b, respectively. Strain, force and strain energy, as well as the EMG activity of the three investigated muscles during walking and running, is depicted in Fig. 4. We found a significant effect of gait speed on maximum AT strain (*p* = 0.043), force (*p* = 0.025) and strain energy (*p* = 0.008, table 3). The post hoc analysis showed significant differences of maximum strain, force and strain energy between walking and fast running (*p* = 0.023, p = 0.018 and p = 0.016, respectively) as well as between slow running and fast running (*p* = 0.023, p = 0.018 and p = 0.016, respectively). The elastic strain energy recoil at the beginning of the stance phase did not show significant differences (*p* = 0.635) between slow and fast running (table 3).

**Figure 4.**
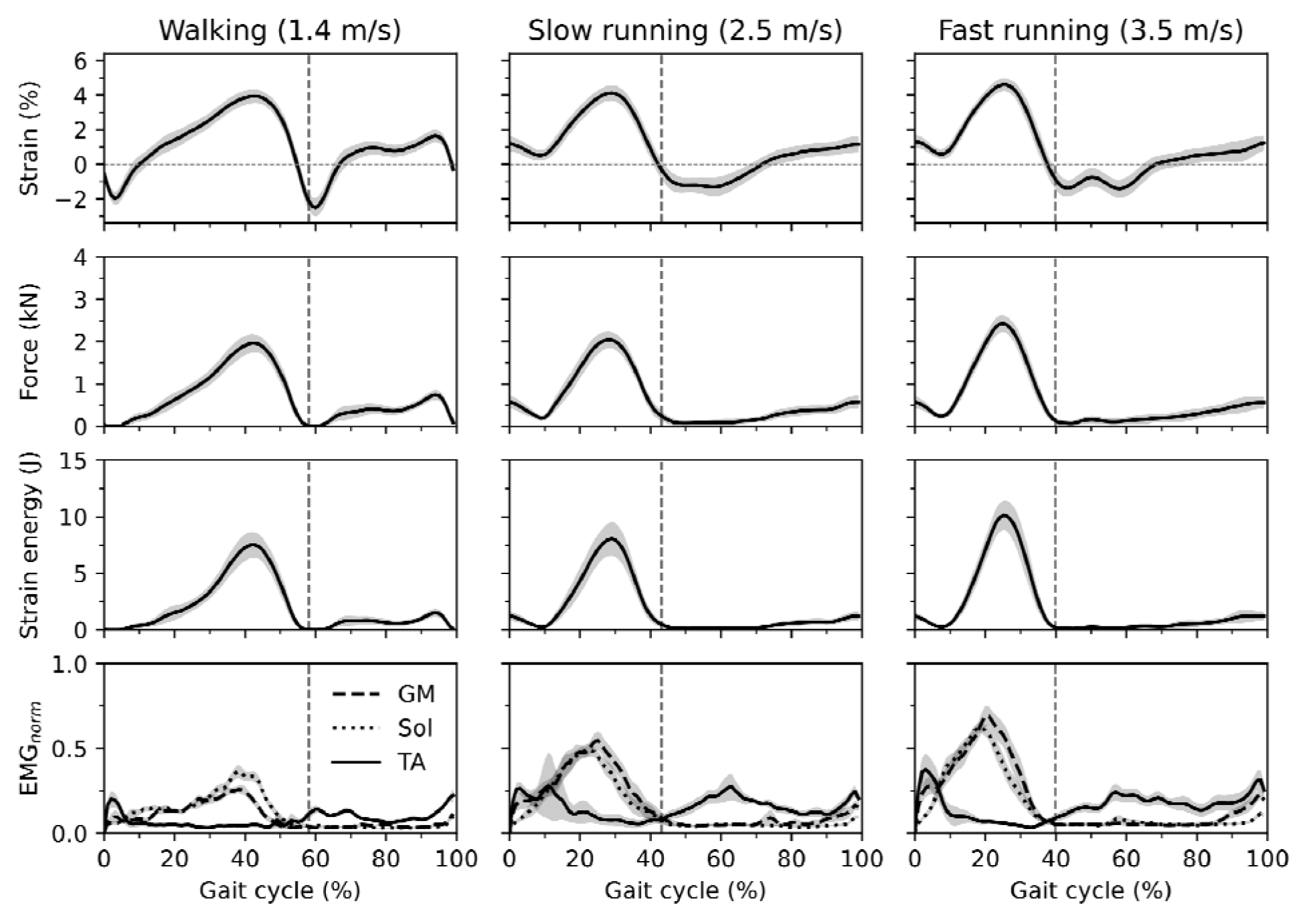
Average values of eleven individuals with ten gait cycle repletion of Achilles tendon (AT) strain, force and strain energy as well as EMG-activity of the gastrocnemius medialis (GM), soleus (Sol) and tibialis anterior (TA) during walking (1.4 m/s), slow running (2.5 m/s) and fast running (3.5 m/s). The gray highlighted area around curves indicates the standard error. The x-axis is normalized to the gait cycle. The gray dashed vertical line separates the contact and swing phase.

**Table 3.** Peak values of strain, force and strain energy of the AT as well as maximum normalized EMG activities of the gastrocnemius medialis (GM_norm_), soleus (Sol_norm_), and tibialis anterior (TA_norm_) during walking and running. Further, elastic strain energy recoil at the beginning of the stance phase of running is presented (average value ± standard deviation).

|  | Walking (1.4 m/s) | Slow running (2.5 m/s) | Fast running (3.5 m/s) |
| --- | --- | --- | --- |
| Strain (%) * | $4.0 \pm 1.2^{\#}$ | $4.5 \pm 1.3^{\#}$ | $4.9 \pm 1.2$ |
| Force (kN) * | $1.989 \pm 0.0684^{\#}$ | $2.284 \pm 0.0536^{\#}$ | $2.556 \pm 0.0540$ |
| Strain energy (J) * | $7.8 \pm 3.7^{\#}$ | $9.5 \pm 4.3^{\#}$ | $11.3 \pm 4.1$ |
| Energy recoil (J) | — | $1.88 \pm 1.10$ | $1.71 \pm 0.65$ |
| $GM_{norm}$ * | $0.31 \pm 0.09^{\# \wedge}$ | $0.68 \pm 0.15$ | $0.8 \pm 0.2$ |
| $Sol_{norm}$ * | $0.38 \pm 0.11^{\# \wedge}$ | $0.61 \pm 0.13^{\#}$ | $0.73 \pm 0.11$ |
| $TA_{norm}$ | $0.27 \pm 0.13$ | $0.54 \pm 0.55$ | $0.5 \pm 0.27$ |

\*: Statistically significant gait effect ( $p < 0.05$ )
#: Statistically significant differences (post-hoc analysis) to fast running ( $p < 0.05$ )
^: Statistically significant differences (post-hoc analysis) to slow running ( $p < 0.05$ )

The maximum TA EMG activity during the stance phase was not significantly different between the three gait speeds (*p* = 0.128). A significant effect of gait speed on maximum GM and Sol EMG-activity was found (*p* < 0.001). The posthoc analysis revealed a significant difference between walking and slow running (*p* < 0.001) and walking and fast running (*p* < 0.001) for both muscles. The post hoc comparisons between slow and fast running revealed a significant difference (*p* = 0.001) only for the Sol muscle (table 3).

## Discussion

In order to assess the mechanical loading and strain energy of the AT during walking and running, we measured AT length using a new in vivo approach, which considers the tendon curve-path shape, taking into account the offset between MTJ and skin surface and the potential movements of the skin relative to calcaneus bone at the AT insertion marker. Further, we developed a novel algorithm to track the MTJ position from the ultrasound image sequences that showed high conformity with manual analysis. The number of video frames we analyzed was for walking in average 1912±411 pictures and for running between 1324 and 1407 pictures for ten gait cycles. The average time needed for the manual tacking of one video was 7.9 hours for walking and 5.4 hours for running. The time for the analysis using our algorithm was considerably reduced to 17 min for walking and 13 min for running, giving evidence for essential improvements of the time-efficiency of the analysis of the MTJ without a decline in quality of the outcome.

The AT morphology features a variable curvature that changes during muscle contraction (Kinugasa et al., 2018, Maganaris et al., 2000). Therefore, the consideration of the AT curved shape is crucial for the investigation of in vivo conditions (Harkness-Armstrong et al., 2020). A reconstruction of the AT curvature using skin markers is a practical and valid approach to address this issue (De Monte et al., 2006, Arampatzis et al., 2008, Fukutani et al., 2014). Our findings indicate that the misalignment of the MTJ and the markers on the skin and skin-to-bone displacements are two important error sources that affect AT length measurements during locomotion when using this approach. The RMS differences between AT length and AT length without MTJ projection, which depicts the contribution of the MTJ projection to the skin surface, were, on average, 2.0 mm (across gait speeds), resulting in maximum AT strain differences of 1.2%. The contribution of the skin-to-bone displacement was lower compared to the MTJ projection. However, the RMSE was, on average, 1 mm, indicating a notable effect on AT length. Taken into consideration the effects of both, i.e., MTJ misalignment and skin-to-bone displacement, the RMS differences increased to 2.5 mm (1.4% strain). During walking and running, the maximum AT strain was, on average, between 4.0 and 4.9%, and therefore, the maximum artifact of 1.4% AT strain resulted in enormous inaccuracies of the AT strain assessment ranging from 28 to 35% during the stance phase of walking and running.

As expected, the maximum strain values in all gait conditions occurred during the stance phase and close to the beginning of the propulsion phase. Shortly after take-off, strain drops rapidly to its minimum value and again increases with a moderate slope until the end of the gait cycle. The maximum strain values were, on average, between 4.0 and 4.9%, corresponding to 51 and 62% of the strain achieved during the MVCs. Repetitive strain of tendons is a crucial mechanical stimulus that regulates cell function and affects the expression of growth factors and the synthesis of matrix proteins (Arnoczky et al., 2002, Lavagnino et al., 2003, Yang et al., 2004), determining tendon plasticity (Arampatzis et al., 2007a, Wang et al., 2013, Bohm et al., 2014). Previous studies examining the effects of submaximal running on AT mechanical properties reported similar AT stiffness between runners and untrained individuals in young and old adults (Karamanidis and Arampatzis, 2005, Arampatzis et al., 2007b) and no adaptive effects of long-time running training on the AT properties (Hansen et al., 2003). Considering that the effective mechanical stimulus for tendon adaptation in terms of strain magnitude ranges between 4.5 and 6.5% strain (Arampatzis et al., 2007a, Bohm et al., 2014, Pizzolato et al., 2019), our findings indicate that submaximal running does not provide sufficient tendon loading for triggering improvements of the AT mechanical properties and may explain the lack of observable effects in the former studies. In both conditions, walking and submaximal running, the applied mechanical load on the AT was likely too low for the initiation of anabolic tendon responses (Pizzolato et al., 2019).

The assessed maximum AT force values in the current study was for walking 2.7 and for running between 3.2 and 3.5 times body weight. These values are significantly lower compared to predictions based on musculoskeletal models, which reported maximum AT forces of 3.9 of body weight during walking (Giddings et al., 2000) and between 5 and 7 of body weight during running (Almonroeder et al., 2013, Lee et al., 2019). These discrepancies might be a result of the limitations of musculoskeletal models, for example, the difficulties in the prediction of activation dynamics using optimization methods (Jinha et al., 2006, Ait-Haddou et al., 2004) for muscle force predictions, which is more evident during rapid movements (Neptune and Kautz, 2001). Further, the missing individual muscle-tendon properties for all involved muscles and the non-consideration of all possible forces that contribute to the resultant ankle joint moment (e.g., ligaments, bones) in the musculoskeletal models enhance the model limitations.

It has been generally accepted that one of the primary roles of the tendon in vertebrates is to store energy during stretching and recoil during shortening (Alexander, 1984, Alexander, 1991). Our results showed an on average maximum energy storage in the AT of 7.8 J during walking, which increased to 9.5 and 11.3 J during slow and fast running. Although the post hoc analysis resulted in a statistically significant difference of strain energy in only fast running compared to walking and slow running, the positive effect of speed on the energy storage and recoil during locomotion is notable and in agreement to earlier studies (Lai et al., 2014; 2015; Monte et al., 2020). During the push-off phase of both walking and running, the whole AT strain energy is returned to the human system (at take-off, the AT strain energy is very close to zero). It is well accepted that this recoil of energy reduces the mechanical work of the plantar flexor muscles (Shadwick, 1990) to accelerate the body’s center of mass in the direction of desired movement (Voigt et al., 1995).

As expected, our results also demonstrated elastic strain energy recoil directly after the touchdown during running. The recoil of strain energy in this initial stance phase (i.e., 70 to 76 ms) was between 1.7 and 1.9 J. These values are 15 to 20% of the maximum AT strain energy during the stance phase and, therefore, might be functionally relevant. We can exclude that this initial strain energy recoil might be dissipated by the muscle contractile elements of the triceps surae muscle because in the investigated running velocities, the fascicles of all three muscles of the triceps surae show an active shortening (Lichtwark et al., 2007, Lai et al., 2014, Bohm et al., 2019). We interpret this finding as the recoil of AT tendon elastic strain energy to the body in this initial phase of stance right after touch down. This phenomenon (i.e., a decrease of AT force and recoil of elastic strain energy after heel contact) has not been mentioned in the literature, and, therefore, the functional consequences for the running task are not known. Komi (1990), using a force transducer which was surgically implanted in the AT, reported a decrease of AT force directly after the touchdown in rearfoot running and suggested an association with the reduction of the TA EMG-activity. As the foot contacts the ground, the TA muscle showed a high EMG activity, indicating an active control of the initial plantar flexion through muscular force. The reduction of the AT force after heel contact decreases the internal resultant ankle joint moment, which supports the function of the TA as a regulator of the ankle joint during the initial part of the stance phase. However, it is difficult to provide a clear explanation of the functional consequences of the resulting AT elastic energy recoil in this initial phase because it is unclear where this energy is returned. A recoil of AT strain energy must not necessarily increase the mechanical work/power at the ankle joint but can, for example, be absorbed by the TA tendinous tissues (Maharaj et al., 2019) or by the elastic structures of the foot arch (Ker et al., 1987, Kelly et al., 2015) and stored as elastic strain energy.

We measured the force-elongation relationship of the AT using MVCs with a slow rate of the force application. The participants completed five trials of isometric ramp contractions, steadily increasing effort from rest to the maximum in ~5 s resulting with an average tendon loading rate of 1.076 ± 0.456 kN/s. Using the individually assessed force-elongation relationship, we calculated the AT force during walking and running. The loading rate of the AT during the stance phase in the investigated conditions ranged from 16.5 to 20.0 kN/s and was significantly higher compared to the MVCs. Tendons, as biomaterials, are viscoelastic, and, therefore, the differences in the loading rate may influence the accuracy of the tendon force assessment during the walking and running trials. However, tendon hysteresis is ~10% (Bennett1 et al., 1986, Pollock and Shadwick, 1994), indicating minor damping components in the tendon properties. Ker et al. (1981) found that loading frequencies from 0.22 to 11 Hz (loading rate from 0.580 to 31.240 kN/s) did not affect the tendon’s youngs modulus (Ker, 1981) and, therefore, we can assume a negligible effect of loading rate on tendon dynamics during physiological activities like walking and running. More recently, Rosario and Roberts (Rosario and Roberts, 2020) investigated the effects of loading rate on tendon strain and predicted a change from slow to fast loading of 0.16% strain on the human AT during running, evidencing a minor effect of loading rate on tendon dynamics. Based on these reports, we can argue that the differences in the loading rate between the MVCs and the investigated locomotor activities did not have significant effects on the AT force assessment.

Further, both the loading and unloading phases of AT force were assessed from the same force-elongation curve during MVC trials and the effect of hysteresis in the unloading phase was not considered. To examine the impact of the hysteresis effect on both AT force and AT strain energy, we assumed a 10% hysteresis, as it is the reported order of magnitude in most studies (Bennett1 et al., 1986, Pollock and Shadwick, 1994, Ker, 1981), and we recalculated the two outputs. For that reason, we determined the two coefficients (a and b) of the Eqn 5 for the unloading phase based on the following constraints: (a) force-elongation curve during the unloading phase will reduce the strain energy by 10%, and (b) both force-elongation curves (loading and unloading) will end in the same point (Fig. 5A). The two constant coefficients of the unloading curve were reached by solving two equations with two unknown variables. The choice of appropriate coefficients in the recalculation of force and strain energy was based on the first derivative of the AT length. In the case that the first derivative was negative, the coefficients of the unloading curve were used; otherwise, the standard force-elongation curve was used (Fig. 5B). The contribution of hysteresis to AT force and strain energy was assessed by means of RMSE, which resulted in a negligible effect up to 46 Newton and 0.2 Joule in forces and energy, respectively (i.e., 1.9% of the maximum AT force or strain energy).

**Figure 5.**
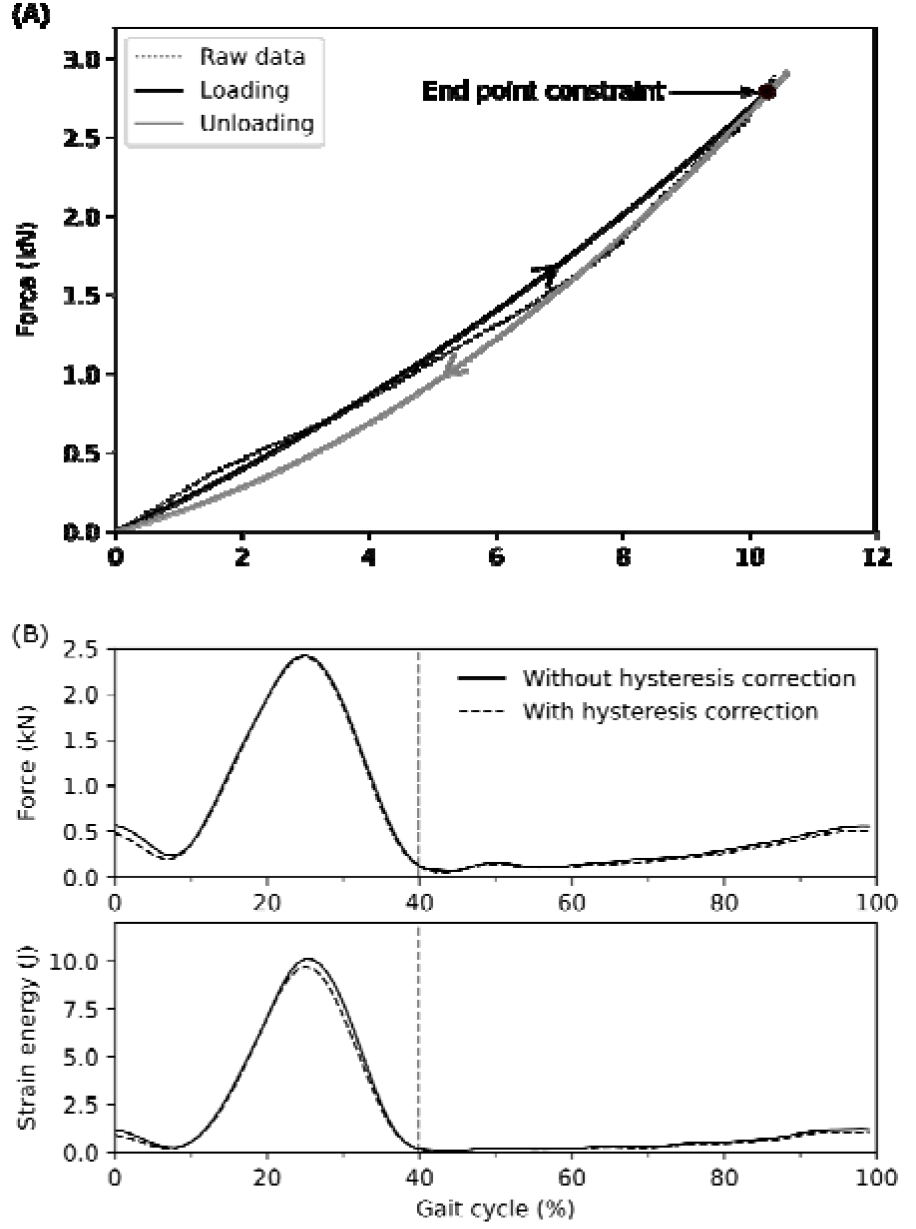
(a) Experimentally assessed Achilles tendon (AT) force-elongation relationship during maximum voluntary contractions (Raw data) and the modeled force-elongation curve during loading and unloading (eleven individuals with ten gait cycle repetition). (b) AT force and AT strain energy with and without the consideration of hysteresis during fast (3.5 m/) running. The x-axis is normalized to the gait cycle. The gray dashed vertical line separates the contact and swing phase.

In conclusion, in this study, we introduced a new in vivo assessment of the AT mechanical loading and strain energy during locomotion, demonstrating that when taking into account the curvature of the AT using skin markers, the projection of the MTJ to the skin misalignment and skin-to-bone displacement are two methodological issues that significantly influence the AT length measurement. We found that the AT mechanical loading during submaximal running is lower than previously reported and too low to initiate an adaptation of tendon mechanical properties, which explains at least partly the reported absence of significant differences in the AT mechanical properties between runners and non-runners. Finally, we provided the first evidence of an elastic strain energy recoil at the beginning of the stance phase of running, which might be functionally relevant for running efficiency.

## Acknowledgements

We thank our colleague Arno Schroll for his help during the data analysis. We acknowledge the Open Access Publication Fund of the HumboldtUniversität zu Berlin.

## Data availability

The datasets generated and analysed during the current study are available at https://figshare.com/s/e4093689f86af36aaaef

